# CDHu40: a novel marker gene set of neuroendocrine prostate cancer (NEPC)

**DOI:** 10.1101/2024.03.28.587205

**Authors:** Sheng Liu, Hye Seung Nam, Ziyu Zeng, Xuehong Deng, Elnaz Pashaei, Yong Zang, Lei Yang, Chenglong Li, Jiaoti Huang, Michael K Wendt, Xin Lu, Rong Huang, Jun Wan

## Abstract

Prostate cancer (PCa) is the most prevalent cancer affecting American men. Castration-resistant prostate cancer (CRPC) can emerge during hormone therapy for PCa, manifesting with elevated serum prostate-specific antigen (PSA) levels, continued disease progression, and/or metastasis to the new sites, resulting in a poor prognosis. A subset of CRPC patients shows a neuroendocrine (NE) phenotype, signifying reduced or no reliance on androgen receptor (AR) signaling and a particularly unfavorable prognosis. In this study, we incorporated computational approaches based on both gene expression profiles and protein-protein interaction (PPI) networks. We identified 500 potential marker genes, which are significantly enriched in cell cycle and neuronal processes. The top 40 candidates, collectively named as CDHu40, demonstrated superior performance in distinguishing NE prostate cancer (NEPC) and non-NEPC samples based on gene expression profiles compared to other published marker sets. Notably, some novel marker genes in CDHu40, absent in the other marker sets, have been reported to be associated with NEPC in the literature, such as DDC, FOLH1, BEX1, MAST1, and CACNA1A. Importantly, elevated CDHu40 scores derived from our predictive model showed a robust correlation with unfavorable survival outcomes in patients, indicating the potential of the CDHu40 score as a promising indicator for predicting the survival prognosis of those patients with the NE phenotype. Motif enrichment analysis on the top candidates suggests that REST and E2F6 may serve as key regulators in the NEPC progression.

**Significance:** our study integrates gene expression variances in multiple NEPC studies and protein-protein interaction network to pinpoint a specific set of NEPC maker genes namely CDHu40. These genes and scores based on their gene expression levels effectively distinguish NEPC samples and underscore the clinical prognostic significance and potential mechanism.

## Introduction

Prostate cancer (PCa) is the most common cancer among American men. With an estimated 288,300 new diagnoses projected for 2023 [1]. Treatments of PCa include surgery, radiotherapy, chemotherapy, hormone therapy, and immunotherapy. However, the development of Castration-resistant prostate cancer (CRPC) during hormone therapy poses a significant challenge. CRPC is characterized by sustained high serum prostate-specific antigen (PSA) levels, ongoing disease progression, and potential metastasis to the new sites, leading to poor prognosis [2–4]. Most CRPC tumors still depend on androgen receptor (AR) signaling, and therefore the use of current androgen-signaling inhibitors (ASI) such as enzalutamide (ENZ) and abiraterone can offer temporary relief from resistance [5]. But a subset of the CRPC patients exhibited neuroendocrine (NE) phenotype with diminished or absent reliance on AR signaling and a dismal prognosis [6–8]. The most lethal subtype of this disease, however, has similar initial symptoms comparable to CRPC, and hence lacks of appropriate unique identification markers. NEPC biopsy samples also often exhibit in mixed histology, posing formidable challenges on accurate diagnosis and appropriate treatments [9].

Thus, developing accurate diagnosis and imaging tools for NEPC is the crucial first step in effectively managing the disease. Numerous studies have attempted to identify the most frequently overlapping markers for NEPC patients including delta-like ligand 3 (DLL3), with high expressions exclusively in CRPC-NE cells [10]. A clinical trial is in progress at Memorial Sloan Kettering Cancer Center to evaluate DLL3 PET imaging in patients with small cell lung cancer (SCLC) and NEPC (NCT04199741). Additional immunohistochemistry (IHC) of canonical NEPC marker genes, such as CHGA, SYP, NCAM1, ENO2, AR, and KLK3, can be used to clinically identify NEPC samples [11], however, IHC of these markers may not always be directly applicable[12]. Given the feasibility of gene expression profiles with the development of next generation sequencing (NGS) technologies, an alternative option is to identify NEPC based on the expression levels and/or other information of gene markers, exploring marker genes beyond the six canonical NEPC markers (Supplemental Table 1).

Despite these efforts, protein-protein interactions (PPIs) have not yet been considered in the selection of NEPC biomarkers in these reports. PPIs are known to form a fundamental network and are integral to almost all biological processes, alteration of which contributes to disease progression. In our study, we incorporated aberrant gene expression and the PPI information by applying the method of using knowledge in network (uKIN)[13] for PPI analysis to explore biomarkers effectively distinguishing NEPC from all of the PCa samples. Our approach facilitates the identification of robust biomarkers associated with NEPC, unveiling novel markers not previously reported as NEPC biomarkers by traditional methods. The top 40 marker genes, denoted as CDHu40, exhibited a remarkable accuracy in predicting NEPC and demonstrated a strong correlation with patient survival. Taken together, our results highlight the potential significance of CDHu40 as a prognostic indicator for NEPC.

## Methods

### Gene expression data sets used in this study

Eight data sets with defined NEPC samples were selected in this study (Table 1). WCM_NEPC_2016 and PRAD_SU2C_2019 were used for the training and validation sets. Three data sets, GSE32967, GSE149091, and GSE59984, were downloaded from the GEO database being used as independent test sets on our model. To our knowledge, samples in Prostate Adenocarcinoma (PRAD) from the TCGA (PRAD_TCGA) were all primary tumors that had barely NEPC cases. So, PRAD_TCGA was used as a negative control to estimate the false positives of the NEPC samples by different marker gene sets. Two more scRNA-seq data sets, Asberry et al 2022[14] and Dong et al 2020[15], were used to test our model at the single cell level.

**Table 1.** Data sets used in this study.

| Data set | Data type | Number of samples |  |  | Reference (PMID or link to the datasets) |
| --- | --- | --- | --- | --- | --- |
|  |  | Total | NEPC | Non-NEPC* |  |
| WCM_NEPC_2016 [6] | Bulk RNA-seq | 49 | 15 | 34 | PMID: 26855148 |
| PRAD_SU2C_2019 [16] | Bulk RNA-seq | 232 | 22 | 210 | PMID: 31061129 |
| GSE32967[17] | Microarray | 22 | 14 | 8 | PMID: 22156612 |
| GSE149091[18, 19] | Bulk RNA-seq | 4 | 1 | 3 | PMID: 32531951, PMID: 32512818 |
| GSE59984[20] | Microarray | 14 | 2 | 12 | PMID: 29757368 |
| PRAD_TCGA | Bulk RNA-seq | 498 | 0 | 498 | https://www.cancer.gov/tcga |
| Asberry et al 2022[14] | scRNA-seq | 4 | 3 | 1 | PMID: 36382181 |
| Dong et al 2020[15] | scRNA-seq | 5 | 4 | 1 | PMID: 33328604 |

\*Non-NEPC represents PCa samples collected in studies which were not recognized as NEPC.

### Identification of differentially expressed genes (DEGs)

Gene expression profiles from the first two datasets in Table 1, WCM_NEPC_2016 [6] and PRAD_SU2C_2019 [16], were retrieved from cBioportal [21, 22]. Limma [23] was used to identify differential expressed genes (DEGs) between NEPC and non-NEPC given the sample information from the above datasets.

### Inference of NEPC associated genes using uKIN

The uKIN [13] was utilized to discover NEPC associated genes strongly associated with the 6 canonical NEPC marker genes: AR, PSA (KLK3), CHGA, SYP, CD56 (NCAM1), and NSE (ENO2), which were used as seeds in the uKIN analysis. The PPI information was retrieved by Physical interaction in the StringDB[24]. DEGs between NEPC and non-NEPC samples were considered as candidates to identify NEPC. The amplitudes of gene expression fold changes (FCs) between NEPC and non-NEPC were added as the weights of nodes/genes for the uKIN analysis.

### Logistic regression model for NEPC classification

Glmnet [[25] was used to generate a logistic regression model to classify NEPC and non-NEPC samples based on marker genes selected. Optimal parameters of the model are estimated using cross validation incorporating elastic net penalty. Beltran et al 2016 [6] and Abida et al. 2019 [16] data were randomly split into training sets including 28 NEPC and 183 non-NEPC samples and test set (9 NEPC and 61 non-NEPC samples). A logistic regression model was built using cross-validation on the training set. The resulting model was tested on the test set and the other three independent GEO test sets (Table 2). The performance of prediction was evaluated based on the area under the ROC curve (AUROC).

**Table 2.**
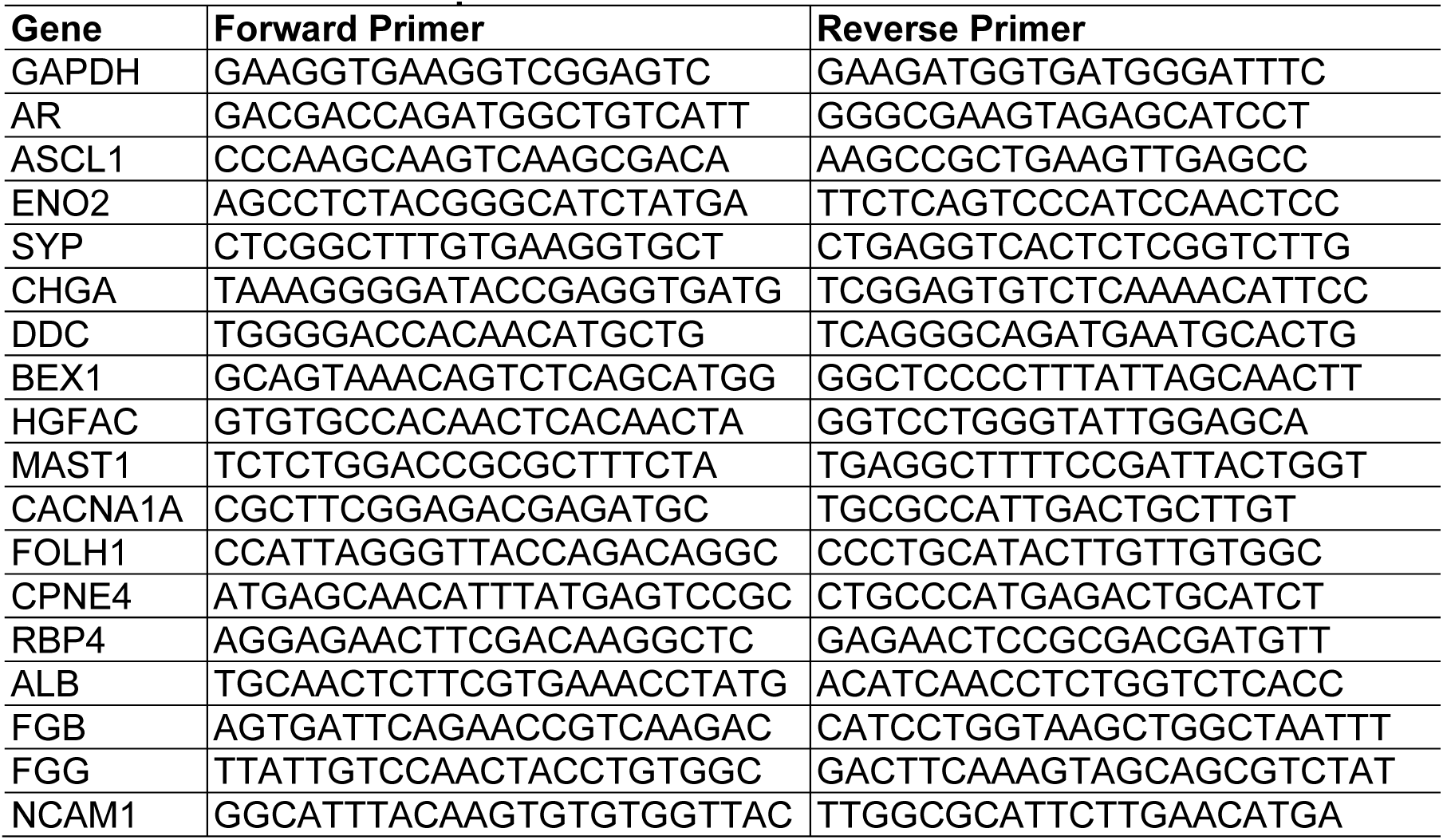
Primers used for qPCR.

### Survival analysis

Survival probability was computed using the function of survfit with Kaplan-Meier (KM) method in the R [26] package survival [27]. Significance of difference in survival times of different groups was determined by log rank test. The KM plots were generated using the ggsurvplot function in the package survminer [28, 29].

### Functional analysis of top candidate genes

The top 500 NEPC candidate marker genes from our results were entered into StringDB [24] for visualization. The functional enrichment analysis was performed using DAVID [30], [31] on top 500 genes with increased and decreased expression in NEPC samples, respectively.

### Cell culture

LNCaP (CRL-1740) and NCI-H660 (CRL-5813) were purchased from American Type Culture Collection (ATCC, atcc.org). The KUCaP13 cell line was obtained from the laboratory of Shusuke Akamatsu at Kyoto University [32]. LNCaP and KUCaP13 cells were cultured in RPMI-1640 medium supplemented with 10% fetal bovine serum (FBS), 1% penicillin-streptomycin (Pen-Strep), and 1% HEPES. NCI-H660 cells were cultured in RPMI-HITES medium containing 5% FBS, 0.005 mg/ml insulin, 0.01 mg/ml transferrin, 30 nM sodium selenite, 10 nM hydrocortisone, 10 nM beta-estradiol, extra 2 mM L-glutamine, and 1% Pen-Strep. LNCaP was passaged at 1:5 ratio every 3-5 days. For NCI-H660 and KUCaP13, half of the medium was refreshed twice a week, and passaged when cell concentration exceeded 1×10^6^ cells/ml. All cell cultures were incubated at 37°C with 5% CO_2_ and assessed for mycoplasma monthly by PCR. All mycoplasma results were negative.

### Gene Expression Detection by qPCR

We further validated the expression levels of 17 selected NEPC markers and GAPDH by qPCR (Table 2) in LNCaP, NCI-H660, and KUCaP13 cells. Total RNA was extracted from cells with EZ-10 Spin Column Animal Total RNA Miniprep Kit (Bio Basic, BS82312), and cDNA was synthesized using All-In-One 5X RT MasterMix (Applied Biological Materials, G592) with 500 ng/μl RNA template, following the respective manufacturer protocols. cDNA was amplified with 2x SYBR Green qPCR Master Mix (Bimake, B21202) and qPCR reaction was run on Bio-Rad CFX Connect system with the following conditions: 95°C for 10 minutes, followed by 40 cycles with 15 seconds at 95°C, and 60 seconds at 60°C, and a final dissociation curve step with 15 seconds at 95°C, 60 seconds at 60°C and 15 seconds at 95°C. Relative fold changes for genes tested were calculated using the ΔΔCt method [33], normalized to GAPDH then compared to LNCaP.

## Results

### Overlap of published marker gene sets

We searched the literature for published NEPC marker gene sets mentioned in the introduction, namely Beltran2016 [6], Tsai2017[34], Bluemn2017[35], Cheng2019[36], Labrecque2019[37], Dong2020[15], Ostano2020[38], Sarkar2023[39], in addition to six most common ones, CHGA, PSA, NCAM1, ENO2, AR, and KLK3, named as NEPC Canonical marker genes (Fig. 1). The majority of genes within distinct marker sets tend to be unique or specific to particular gene expression dataset employed for identification. The maximum pairwise overlap was observed for 11 genes between Beltran2016 and Tsai2017. Even the six canonical NEPC markers were rediscovered in certain marker gene sets but not universally across all of them. This lack of consensus in NEPC markers suggests the intricate nature of biological processes associated with NEPC progression and potential biases when simply comparing gene expression differences for different datasets.

**Figure 1.**
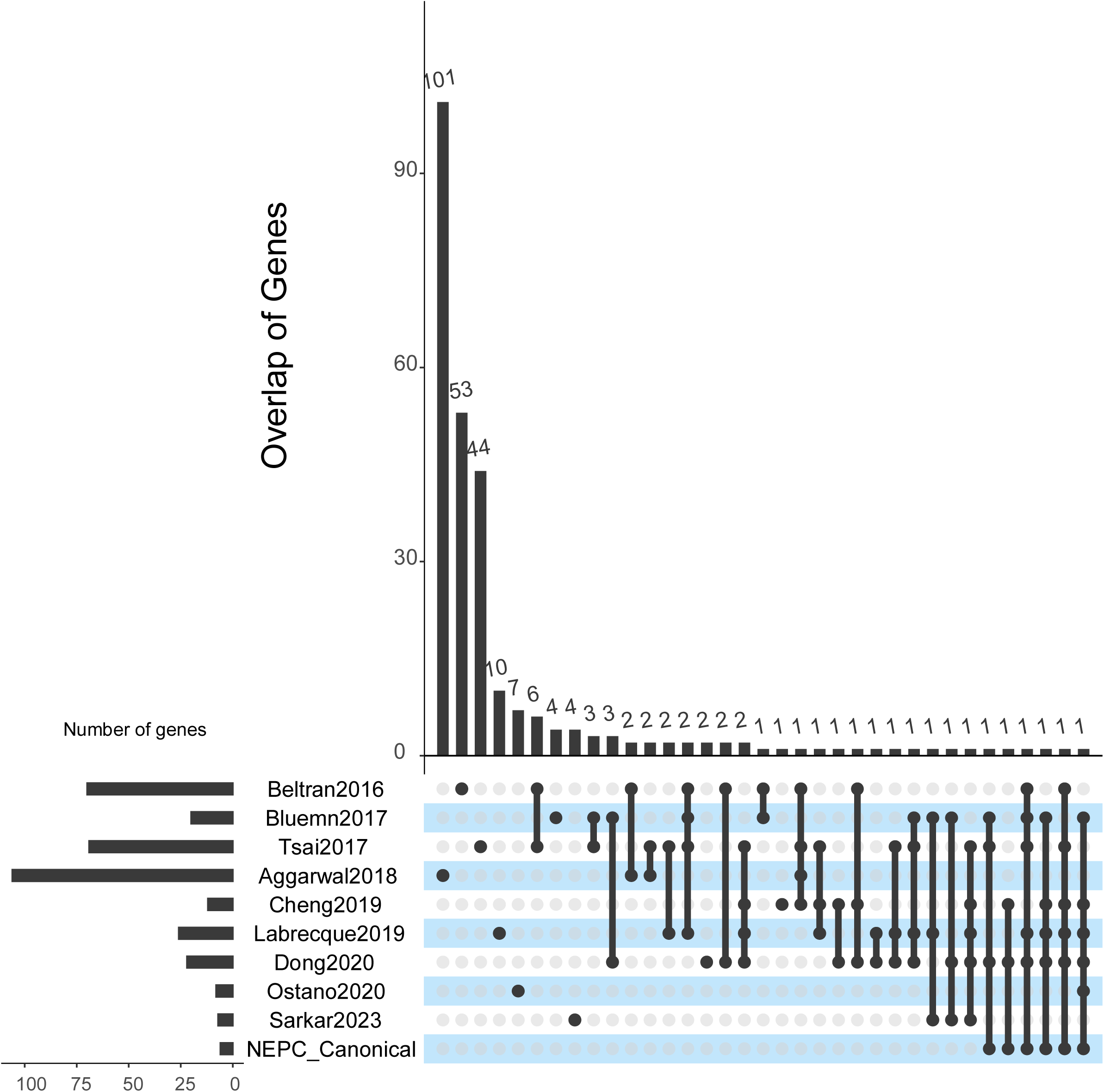
Overlap of NEPC marker genes published by selected literatures.

### Integration of uKIN identifies higher confidence NEPC-related genes

To identify NEPC biomarkers combining different gene expression datasets while considering protein-protein interactions (PPIs), we utilized uKIN to pinpoint potential biomarker genes (Fig. 2). Specifically, we performed differential analysis between 15 NEPC and 24 non-NEPC samples in Beltran et al 2016 data and between 22 NEPC and 210 non-NEPC samples in Abida et al 2019 data, respectively. Then we applied uKIN taking the six canonical NEPC markers as seeds with the support of PPI networks and using FC amplitudes of gene expression between NEPC and non-NEPC as weights for the new information added into the network to rank the NEPC relatedness of the genes. The top genes from the uKIN were considered potential NEPC biomarkers. Next, we derived a model based on selected top genes and further performed a functional analysis of top genes from the uKIN.

**Figure 2.**
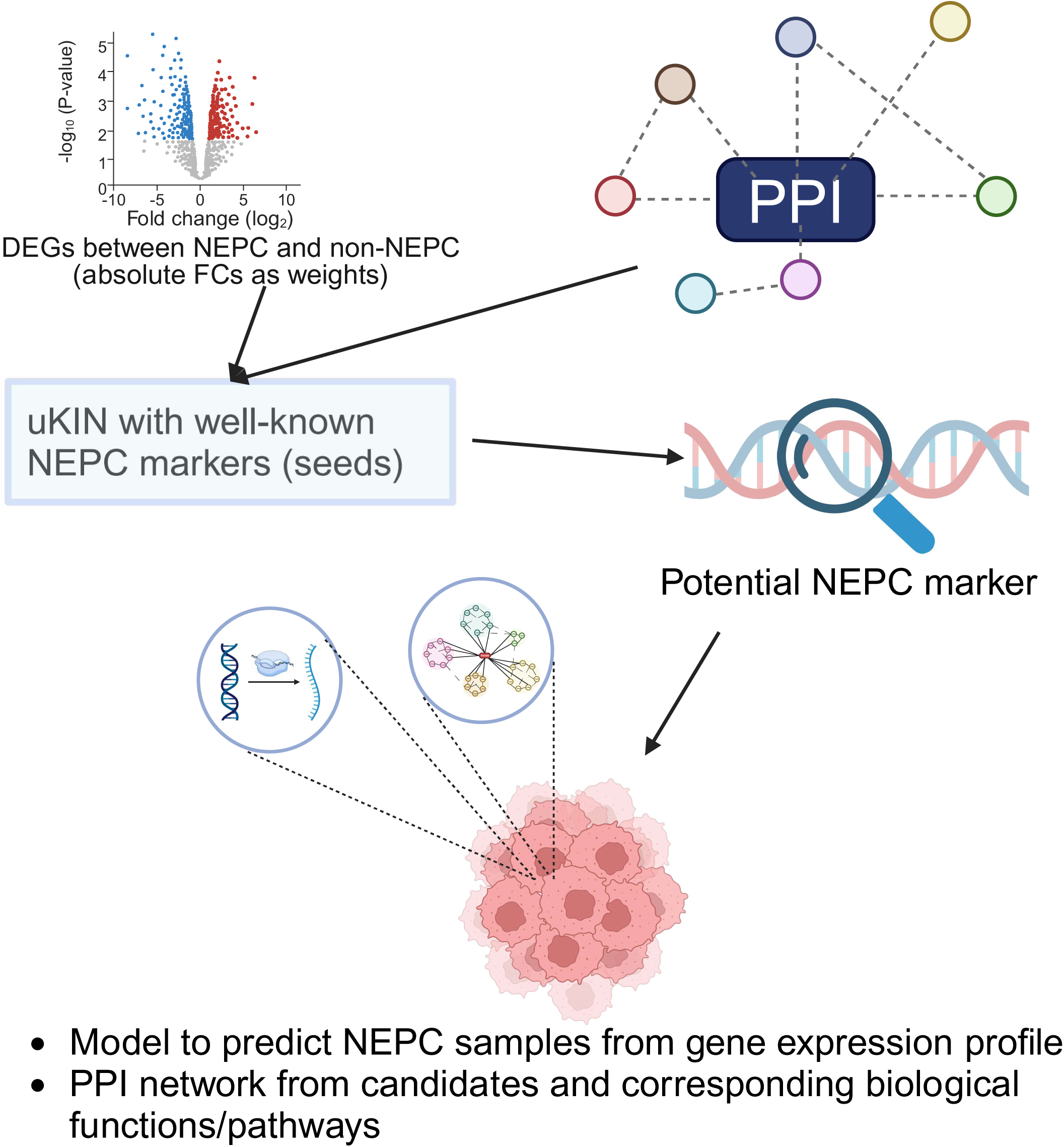
Flow chart of our approach. PPI: Protein-Protein Interaction. DEG: Differential expression genes. FC: Fold change. uKIN: using knowledge in Network.

### Selection of top performing gene sets

To assess the effectiveness of identified top potential biomarker genes in predicting the NEPC phenotype, we applied elastic net logistic regression to different numbers of top ranked genes identified by our approach, including top 10, 20, 30, and up to top 100 genes, respectively, aiming to select the optimal parameters that yield a highly regularized model with elastic net penalty. The 10-fold cross-validation and 10 repetitions were taken using the gene expression profiles in the training set to ensure that the cross-validated error falls within one standard error of the minimum.

Taking into account the balance between the performance and the quantity of marker genes involved, we chose the top 40 candidate genes, namely CDHu40, as the gene set for NEPC biomarkers. The CDHu40 outperformed most of other NEPC marker gene sets listed here, except Beltran2016 and Dong2020 (Fig. 3A), in terms of AUROC scores on both test datasets and other independent GEO datasets collected in the study.

**Figure 3.**
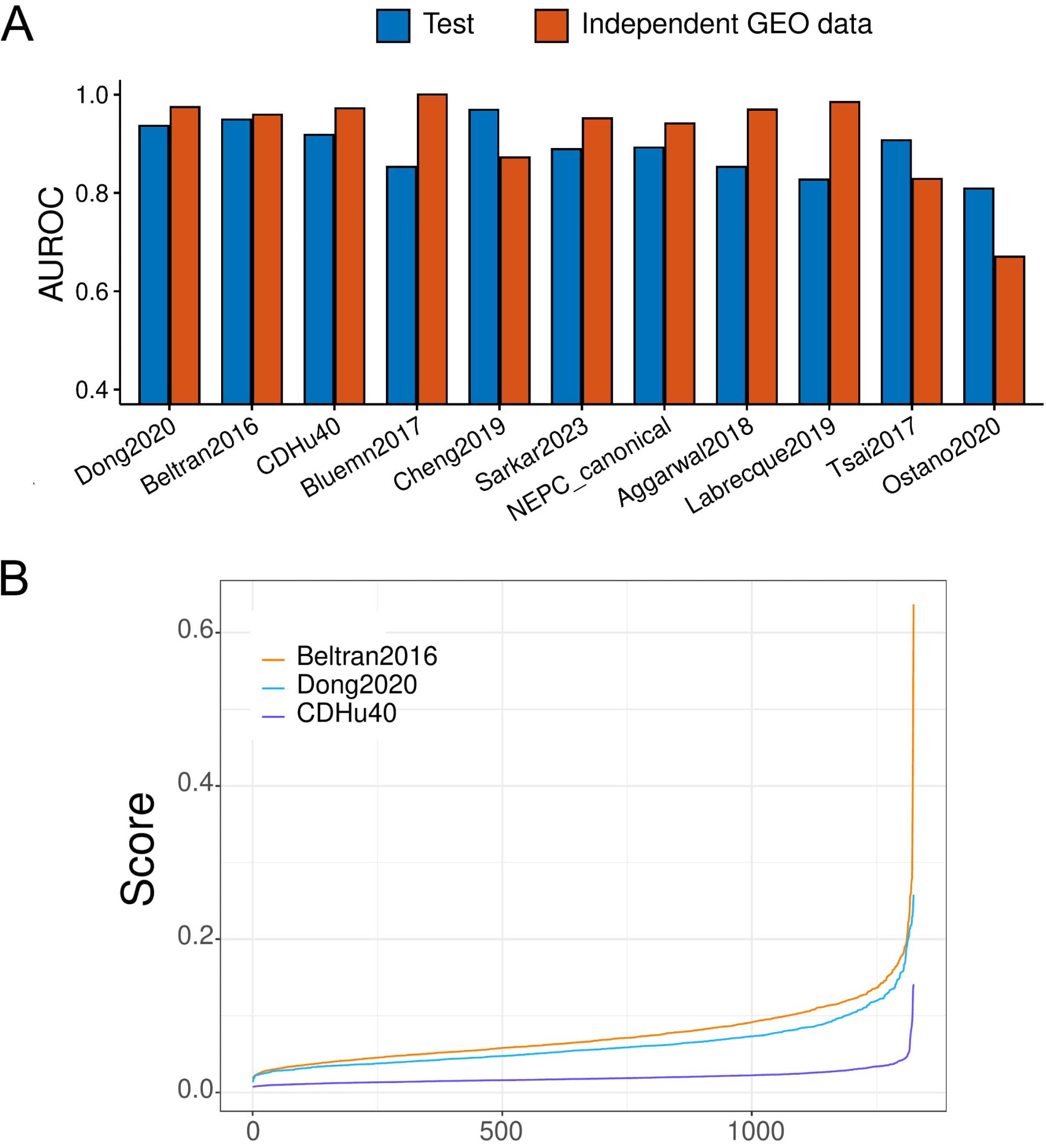
Performance of CDHu40 and other published NEPC marker gene sets. (A) Bar plot of AUROC for each set. Gene sets were sorted by the average values of AUROC. (B) NEPC scores of PRAD-TCGA samples estimated by Beltran2016, Dong2020, and CDHu40, respectively.

CDHu40 score was defined as the NEPC prediction probability using elastic net logistic regression model. Additionally, we computed NEPC scores using two other marker gene sets, Beltran2016 and Dong2020, respectively, since both of them also exhibited superior performance on two test sets in the study. We calculated these scores utilizing gene expression data of PRAD_TCGA (Fig. 3B). In general, CDHu40 scores were notably lower compared to the scores generated based on Dong2020 and Beltran2016 gene sets. Some samples exhibited higher scores according to Beltran2016. Given that the PRAD_TCGA samples were from primary PCa tumors that lack a NE phenotype, this suggests that CDHu40 provides a more accurate representation of the NEPC phenotype compared to Dong2020 and Beltran2016, particularly in terms of minimizing false positive rates.

Except genes of CDHu40 recovered by other marker gene sets previously mentioned (Fig. 4A), more than a quarter (11 out of 40) of CDHu40 genes were absent from any of these sets (highlighted in red in Fig. 4A), although some of them have been investigated in independent studies. For instance, DDC was reported as a neuroendocrine marker in various human tumors originating from NE cells [40]PMID: 3383200. Prostate-specific membrane antigen (PSMA), often overexpressed in most prostate adenocarcinoma (AdPC) cells, serves as a marker for PC and becomes a target for molecular imaging.

**Figure 4.**
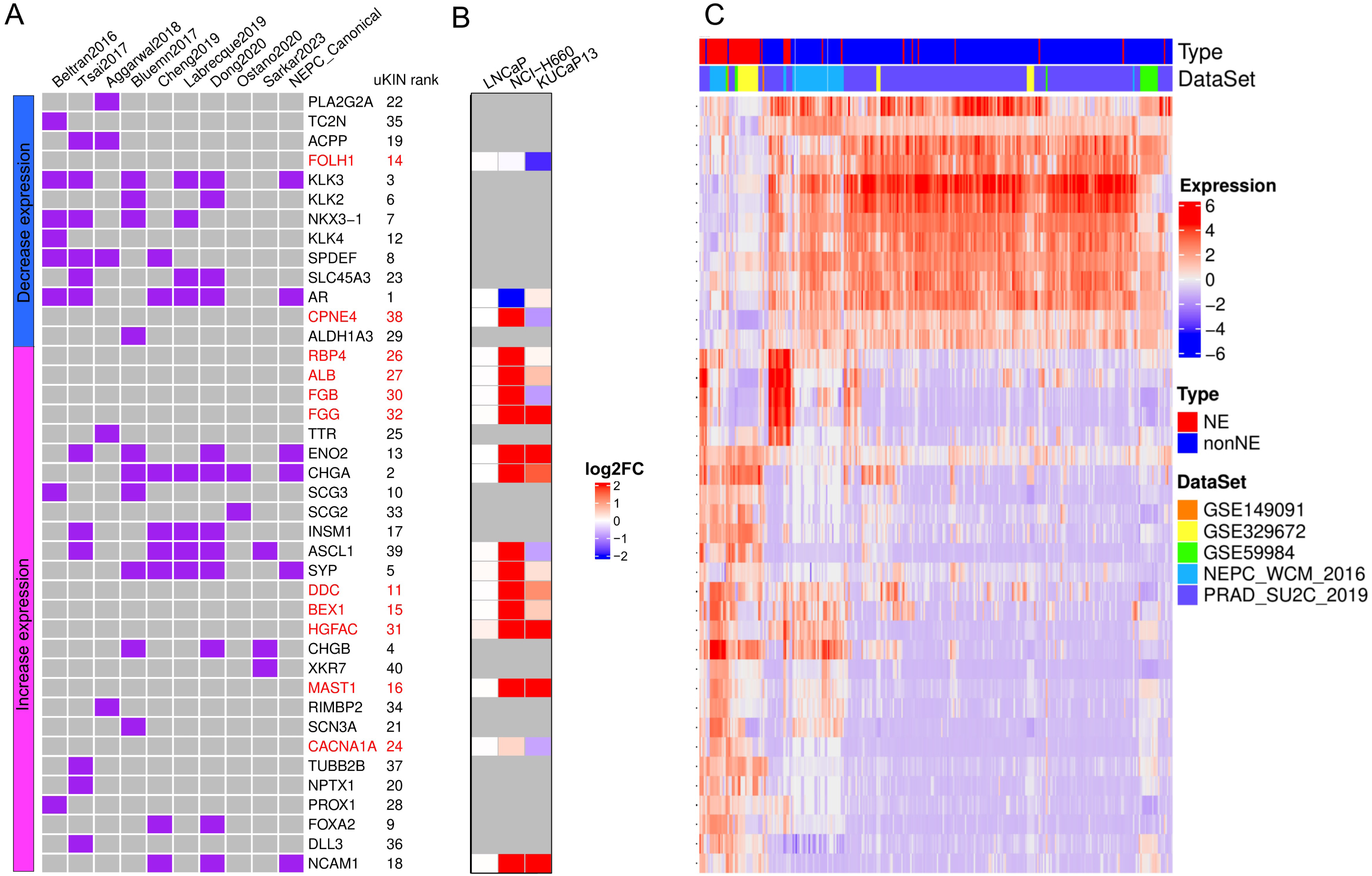
CDHu40 genes identified. (A) Overlap of CDHu40 and other marker gene sets in the literature. The left bar shows that marker genes were either down- (blue) or up- (purple) regulated in NEPC samples as we identified. Genes highlighted by red color were absent from all other published marker genes compared here. (B) qPRC validation on 17 selected marker genes in three PCa cell lines, LNCaP, NCI-H660, and KUCaP13. (C) Expression profile of CDHu40 genes obtained by different data sets.

The down-regulation of the PSMA gene (FOLH1) in NEPC samples rendered it a marker capable of distinguishing NEPC from AdPC and this suppressed expression of PSMA gene in NEPC results in failure of NEPC identification using PSMA-targeting imaging. [41]. BEX1 was recognized for its involvement in the tumorigenesis of NE-specific tumors [42] [43] [44]. MAST1 played a role in modulating neuronal differentiation and cell cycle exit through P27 in neuroblastoma cells [45]. Notably, mutations in CACNA1A were linked to neuroendocrine dysregulation. [46]. ALB was identified as an independent risk factor for lymph node metastasis (LNM) in gastric NE tumor (G-NET) patients.[47]. FGB and FGG, both up-regulated in duodenopancreatic NE tumors (DPNETs), signified a de-differentiation process in DPNET patients with poor outcomes [48]. Additionally, the fusion of CPNE4 and ACAD11 was identified in NE samples [49]. The qPCR experiments in three PCa cell lines, LNCaP, NCI-H660, and KUCaP13 were conducted to test the expression levels of these novel marker genes in addition to several well-known NEPC markers (Fig. 4B). The expression changes of most identified novel marker genes aligned with the predicted either up or down regulation in at least one of the two NEPC cell lines. Collectively, they indicate that many unique genes in CDHu40 are strongly associated with the NEPC phenotype and warrant further in-depth understanding and investigations.

Fig 4C showcases the expression patterns of these CDHu40 genes by two-way clustering across various independent datasets with documented NEPC information that were generated by independent groups and collected for our study. Discernible are two major groups of genes, either down-regulated in NEPC samples, exemplified by AR, KLK3, FOLH1, etc., or up-regulated in NEPC samples, e.g., CHGA, ENO2, SYP, DDC, BEX1, HGFAC, and others. However, several subsets of non-NEPC samples were observed with higher expression of RBP4, ALB, FGB, FGG, and TTR, or DDC, BEX1, HGFAC, and CHGB (Fig 4C), which typically show augmented expression levels in the majority of NEPC samples. It suggests potential subtypes of certain documented non-NEPC samples that might come with some NEPC features or were progressing toward NEPC. Such information may bring new insights into the molecular mechanisms of the development of NE phenotype from PCa.

### Validation of CDHu40 score in multiple PCa datasets

We re-analyzed two sets of previously published scRNA-seq data[14, 15] and subsequently applied the CDHu40 score at the single cell level to distinguish NEPC cells from others. Significant enrichments of cells with higher CDHu40 scores were observed in the clusters 5 and 17, as well as AR-cells in cluster 2 (Fig. 5A) at Day 14 subjected to enzalutamide (ENZ) treatment inducing neuroendocrine differentiation (NED), consistent with the results reported and the phenotype changes observed in the experiments [14].

**Figure 5.**
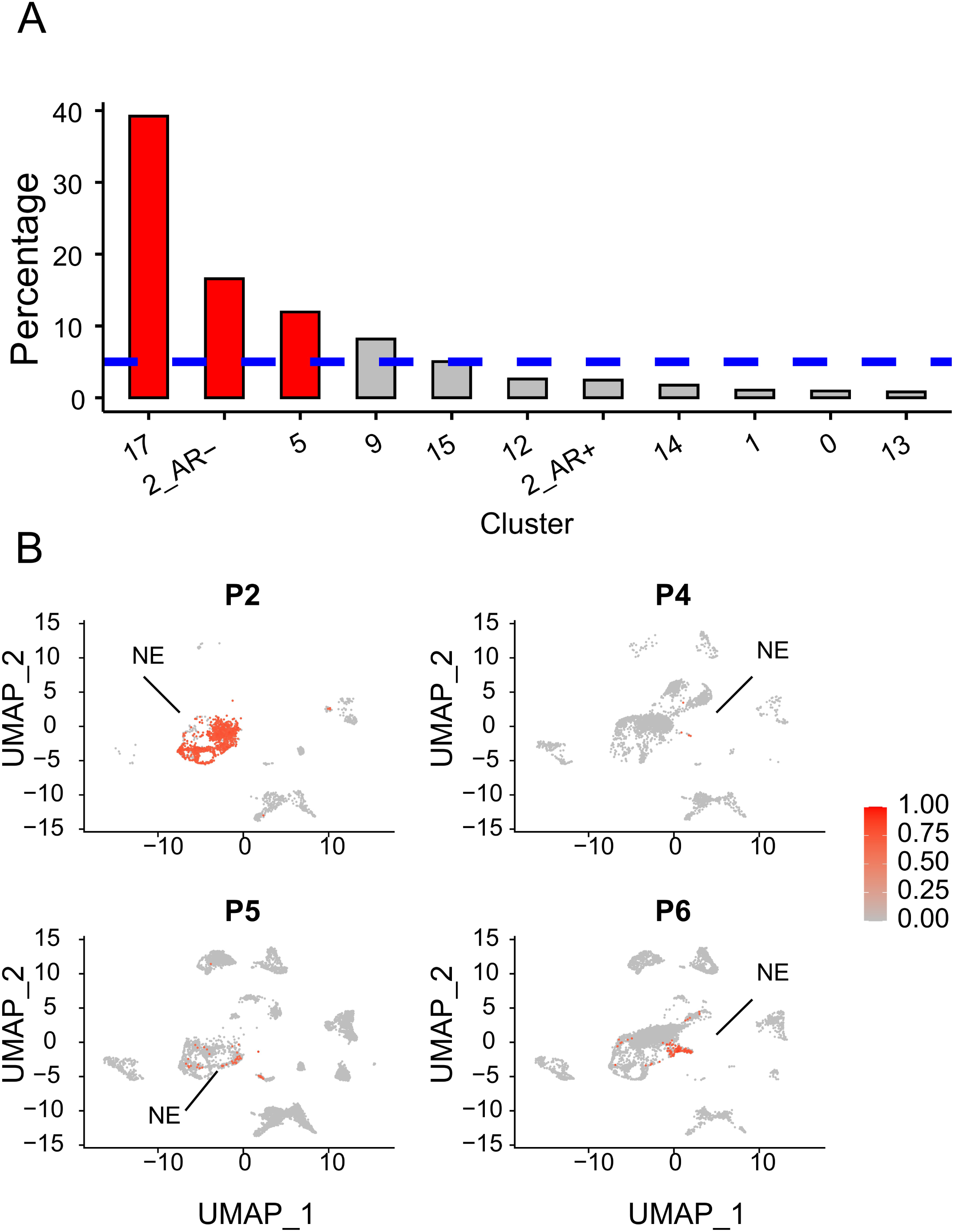
Cells with higher CDHu40 scores from scRNA-seq datasets. (A) Percentage of cells with higher CDHu40 scores in each cluster recognized by Asberry et al 2022 at Day 14 after neuroendocrine differentiation (NED). The red bars represent the significantly (p < 0.05) higher ratio, whereas the blue dashed line is the average percentage of high-CDHu40 score in all cells in the sample. (B) Cells with higher CDHu40 scores were marked in red given the patient samples with the scRNA-seq by Dong et al 2020.

Dong et al [15] published scRNA-seq datasets based on samples from CRPC patients characterizing the tumor cell diversity in 2020. Among these patients, four were clinically determined to have undergone NED. NE cells showed significantly higher CDHu40 scores compared to non-NE cells (Fig 5B), underscoring the robustness of CDHu40 score even at the single cell level.

### Strong correlation between CDHu40 and survival data

Taking the dataset of PRAD_SU2C_2019 [16] as another example, we evaluated scores estimated by the CDHu40, Dong2020, and Beltran2016, respectively, across all NE and non-NE samples (Figs. 6A-C) to examine these scores with the clinical diagnoses. CDHu40 scores displayed a significant increase in NE samples in comparison to non-NE samples, aligning consistently with the NE phenotype diagnosed by clinicians. More strikingly, patients with higher CDHu40 scores experienced poor overall survival (OS) times with statistical significance (*p* = 0.03) using the log rank test (Fig. 6D). But no notable difference was observed for patients classified by higher or lower scores based on Beltran2016 and Dong2020 (Figs. 6E&F) or generated by other marker sets and (Supplemental Fig. 1).

**Figure 6.**
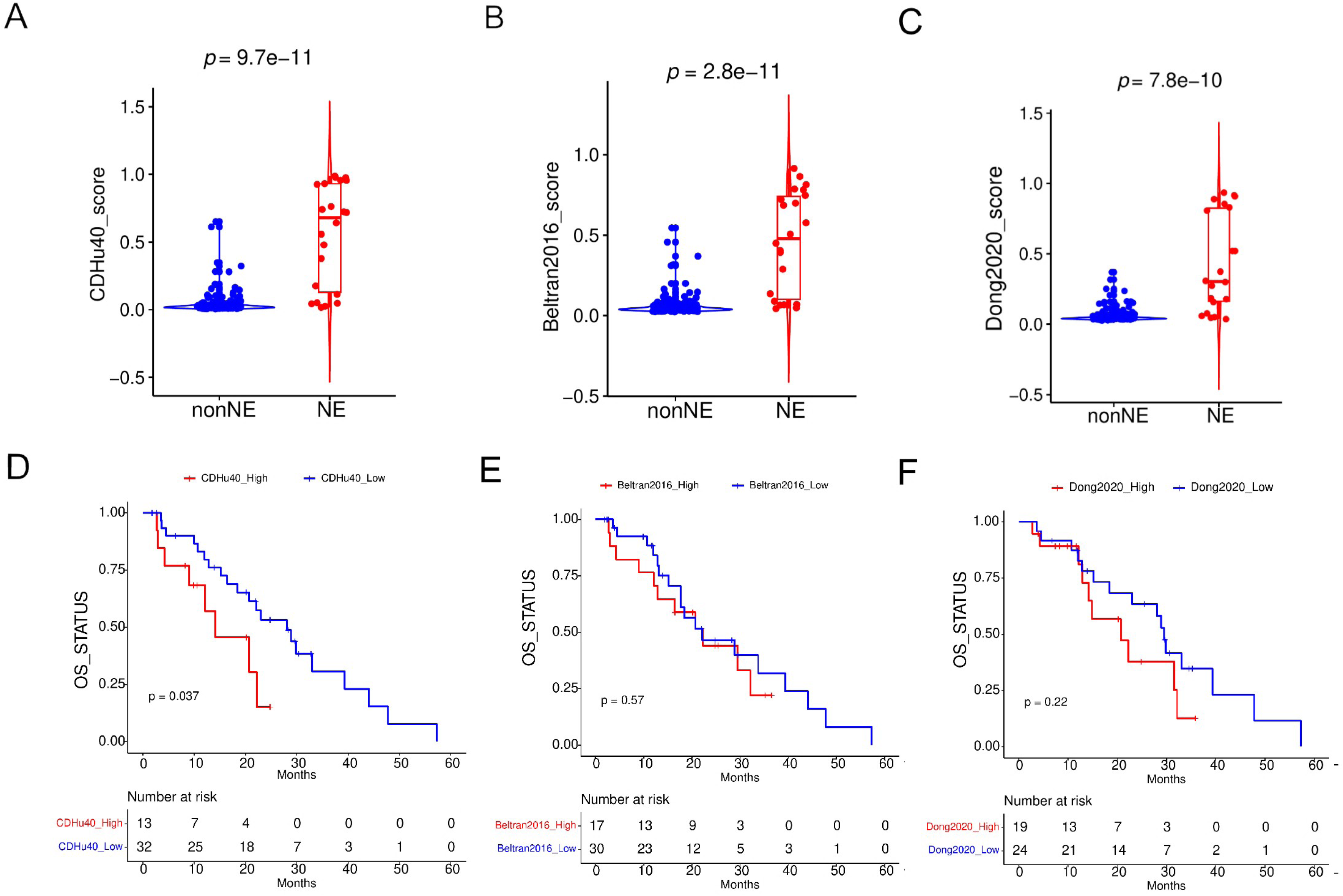
Examination of CDHu40 score on PRAD_SU2C_2019 samples. NEPC scores on clinical NE and non-NE samples were calculated by (A) CDHu40, (B) Beltran2016, and (C) Dong2020. Corresponding survival differences were evaluated based on scores by (D) CDHu40, (E) Beltran2016, and (F) Dong2020, respectively.

Similar trends were evident when we tested the PRAD-TCGA datasets (Supplemental Fig. 2). Patients with augmented CDHu40 scores encountered significantly (log rank test *p* = 3.0×10^-4^) shorter disease-free survival (DFS) times than those with lower CDHu40 scores (Supplemental Fig. 2A). No remarkable differences were noted between higher and lower scores identified by either Dong2020 (Supplemental Fig. 2B) or Beltran2016 (Supplemental Fig. 2C). These findings indicate the potential of the CDHu40 score as a promising prognostic indicator of patients with the NE phenotype.

### Functional analysis on top 500 candidate genes

We performed biologically functional enrichment analysis by using DAVID [30, 31] on the genes that were up- and down-regulated, respectively, in NEPC samples among the top 500 candidates identified in the study. The analysis revealed some biological processes significantly enriched in up-regulated genes, such as the generation of neurons, neurogenesis, regulation of neuron differentiation, and cell cycle (Fig. 7A), in accordance with NE features observed. Interestingly, genes activated in NEPC samples enriched for protein activation cascade may suggest their potential contributions to neurogenesis or cell cycle processes.

**Figure 7.**
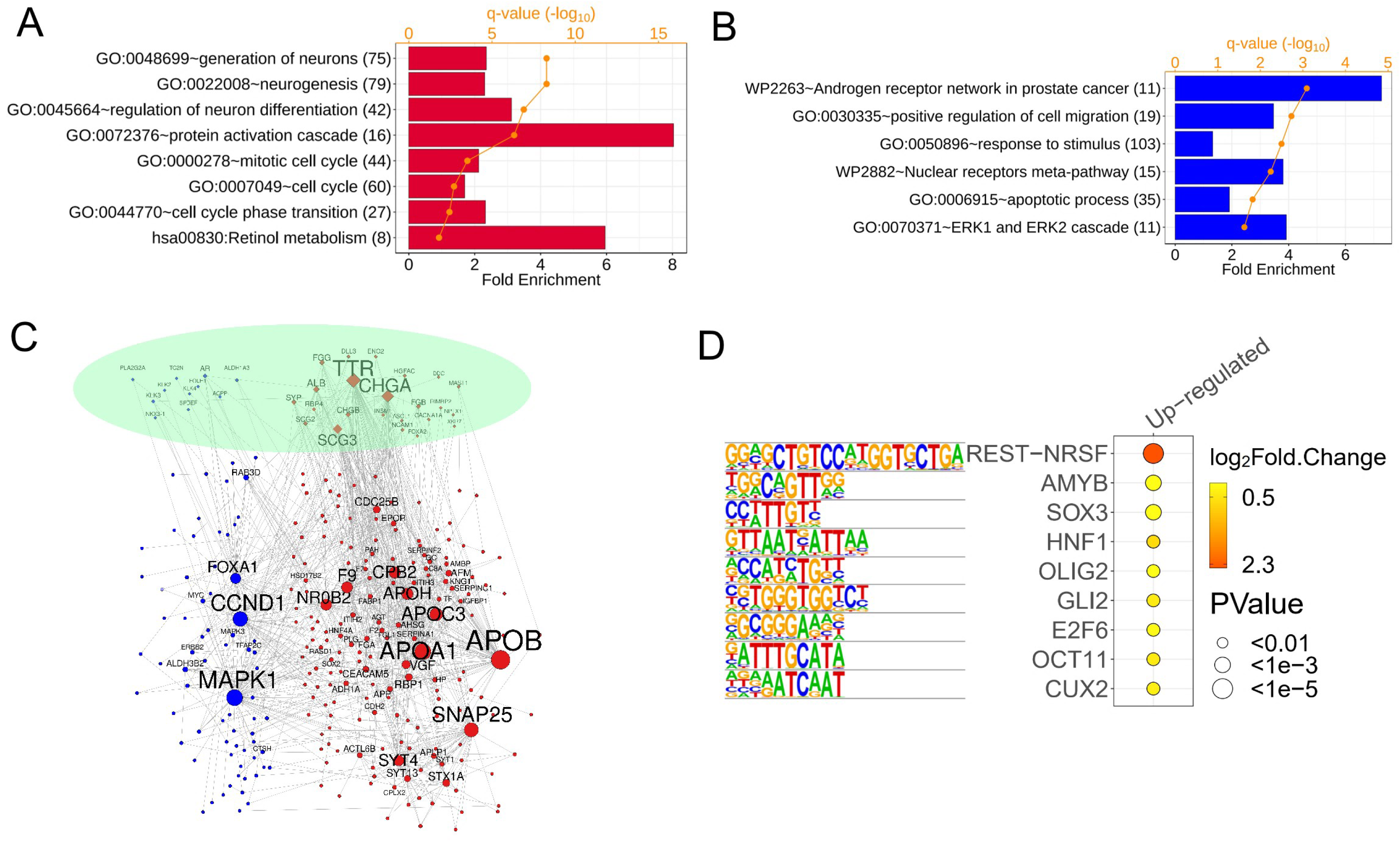
Top 500 genes identified by our methods. Gene ontology (GO) and KEGG pathway enriched in (A) up-regulated genes and (B) down-regulated genes. (C) PPI network of top 500 genes. Red and blue nodes are genes with elevated and lower expression levels, respectively, in NEPC samples. Yellow diamond nodes are CDHu40 genes. (D) Motifs enriched in the regions from upstream (2kb) to downstream (500bp) of 330 up-regulated candidate genes.

Conversely, an enrichment of the AR network in PCa was observed in down-regulated genes (Fig. 7B), corresponding to the inhibition of AR expressions in NE samples. Additionally, apoptotic processes and cell migration were enriched in down-regulated genes, suggesting repressed cell migration might affect or even lead to a shift toward a more vital NE status.

Fig. 7C exhibits the PPI network for the top 500 candidate genes. Among 40 CDHu40 genes, 11 genes serve as hub genes, being connected with at least 5 other genes in the network. This indicates the importance of CDHu40 in establishing connections among genes leading to the featured functions of NEPC. Other hub genes such as CCND1 [50] and CDC25B [51] are associated with the cell cycle. CDC25B induces cellular senescence and correlates with tumor suppression in a p53-dependent manner. Additionally, APOA1 was identified to be upregulated in normal PC. Augmented APOA1 reflects its potential role in driving therapeutic resistance and disease progression by reprogramming the lipid metabolic network of tumor cells [52]; APOB, APOC3, and APOH may function like APOA1. Other highlighted gene was SYT4, which is a well-characterized marker for NE tumors, [53] [54]. NR0B2 is a novel androgen receptor co-repressor in mouse Sertoli cells [55]. MAPK1 plays a role in the activation of Erk1/2- mitogen-activated protein kinases (MAPK) signal transduction pathway in SCLC [56], which has NE features. FOXA1 inhibits prostate cancer neuroendocrine differentiation [57].

We investigated the consensus sequences in the region spanning upstream 2kb to downstream 500bp of up-regulated genes and down-regulated genes, respectively, in the top 500 gene candidates (Fig. 7D). The repressor element-1 (RE-1) silencing transcription factor (REST) motif was observed to be enriched in the up-regulated genes, including targets CHGA, CHGB, and SYP based on the motif analysis. The known functions of REST involve the repression of neural genes and the negative regulation of neurogenesis. Up-regulation of REST target genes in NEPC samples indicates a weakening or absence of the repression role of REST, in turn activating or contributing to the NE features. Another noteworthy finding is the involvement of E2F6, a member of the E2F family, activated in the NEPC samples. E2F6 plays a critical role during the G1/S transition in the mammalian cell cycle. This suggests potential dysfunction of the cell cycle in NEPC through the activity of E2F6 and its target genes. Interestingly, no significantly enriched motifs were observed in down-regulated genes among the top 500 candidates.

## Discussion

Prostate cancer (PCa) stands as the most prevalent cancer among men in the United States[1]. Hormone treatment is the frontline treatment regimen for prostate cancer patients due to the disease sensitivity towards androgen. All existing therapies for prostate cancer, particularly the next-generation ASI drugs, lead to the development of particular subtype of prostate cancer that exhibits NE-like phenotype that no longer responsive to any type of antiandrogen treatment [58]. Not only NEPC is treatment-resistant, but its complex genetic heterogeneity also contributes to misdiagnosis and challenges in recognition.

Due to the lack of biopsy samples of metastatic NEPC, poor characterization of the disease is still prevalent [9]. One of the hurdles of disease identification is the consequent under sampling of mixed histology of the NEPC samples. Genotypic and phenotypic evaluation only represents small lesions of the actual diverse genetic profile of the disease. Several studies have shown potential diagnosis markers for NEPC such as CGA, NCAM1, SYP, and NSE, however, their expressions do not always coincide among patients. To address this biopsy barrier, we established potential diagnostic and prognostic markers for NEPC patients.

Here we proposed a novel integration method incorporating differential gene expression analysis between NEPC and non-NEPC samples as well as the uKIN algorithm based on the PPI network starting with several well-known NEPC biomarkers. The approach effectively generates a list of candidate biomarker genes for NEPC. Our analysis of the top 500 candidates revealed enrichment in neural-related features and cell cycle process enriched in genes up-regulated in NEPC, along with repression in the AR network in NEPC. The PPI network for these top 500 genes identified hub genes associated with the cell cycle and progression of NED. Additionally, motifs of REST and E2F6 were enriched in promoter regions of these top candidates, suggesting their involvement in the generation of NE features and cell cycle regulation.

We specifically selected the top 40 candidate genes, termed CDHu40, which includes some novel targets not reported in other NEPC marker sets. The CDHu40 gene set exhibited functional relations with NE features, providing further insights into the underlying NED mechanisms. Particularly, CDHu40 genes demonstrated robust and efficient performance in predicting NEPC samples using both bulk mRNA expression and single cell expression data compared with other published marker gene sets. More importantly, the CDHu40 score emerges as a better diagnostic marker for NEPC and a reliable prognostic marker for NEPC patients.

Nevertheless, due to the heterogeneity of NEPC, most of the time, the identified markers will not reflect the clinical identification of NEPC/NE-CRPC. For example, variations in CDHu40 gene expression profiles were observed across diverse datasets (Fig. 4B). Notably, distinct subsets of non-NEPC samples were noted with elevated expressions of either RBP4, ALB, FGB, FGG, and TTR, or DDC, BEX1, HGFAC, and CHGB (Fig. 4B). These CDHu40 marker genes typically exhibit heightened expression levels in the majority of NEPC samples. This observation suggests potential subtypes within documented non-NEPC samples that may harbor some NEPC features or could be progressing toward NEPC. Collectively, these insights contribute to improving diagnosis and imaging options for NEPC patients.

## Authors’ Contributions

S. Liu: Conceptualization, resources, data curation, software, formal analysis, investigation, visualization, methodology, writing-original draft, writing-review and editing. H.S. Nam: Conceptualization, resources, data curation, validation, investigation, visualization, writing-review and editing. Z. Zeng: Validation, investigation, methodology, writing-review and editing. X. Deng: Conceptualization, resources, data curation, validation, investigation, visualization, project administration, writing-review and editing. E. Pashaei: formal analysis, investigation, methodology, writing-review and editing. Y. Zang: Conceptualization, software, formal analysis, funding acquisition, investigation, methodology, project administration, writing-review and editing. L. Yang: Conceptualization, funding acquisition, investigation, methodology, writing-original draft, project administration, writing-review and editing. C. Li: Conceptualization, investigation, funding acquisition, project administration, writing-review and editing. J Huang: Conceptualization, resources, investigation, funding acquisition, project administration, writing-review and editing. M.K. Wendt: Funding acquisition, investigation, visualization, writing-review and editing. X. Lu: Conceptualization, resources, data curation, validation, formal analysis, investigation, visualization, methodology, funding acquisition, project administration, writing-review and editing. R. Huang: Conceptualization, resources, data curation, funding acquisition, investigation, project administration, writing-review and editing. J. Wan: Conceptualization, resources, data curation, investigation, visualization, methodology, software, formal analysis, writing-original draft, writing-review and editing, funding acquisition, supervision & project administration.

## Acknowledgments

The authors express gratitude to Dr. Chang-Deng Hu for his valuable insights and support of the project. Regrettably, Dr. Hu succumbed to a rare and highly aggressive cancer in September 2022. In honor of his remarkable contributions to this work and his lifetime dedication to groundbreaking cancer research in biochemistry and pharmacology, we have designated the marker gene set as CDHu40.

This work was partially supported by the grants from the National Institutes of Health (NIH, R01CA248033 to XL), the U.S. Army Medical Research Acquisition Activity, Prostate Cancer Research Program (W81XWH-13-1-0398 to CDH, W81XWH-16-1-0394 to CDH, and W81XWH-22-1-0332 to RH), Indiana University Simon Comprehensive Cancer Center (IUSCCC, NIH/NCI P30CA082709), The Near-Miss Initiative at IUSCCC to JW, the Showalter Scholar program funded by Ralph W. and Grace M. Showalter Research Trust Fund to JW and LY, and additional support from the Walther Cancer Foundations. We are also grateful for the insightful discussions with previous lab members in Dr. Chang-Deng Hu’s lab, including Drs. Andrew Michael Asberry, Jake L Owens, and Elena Beketova.

---

**Supplemental Table 1.**
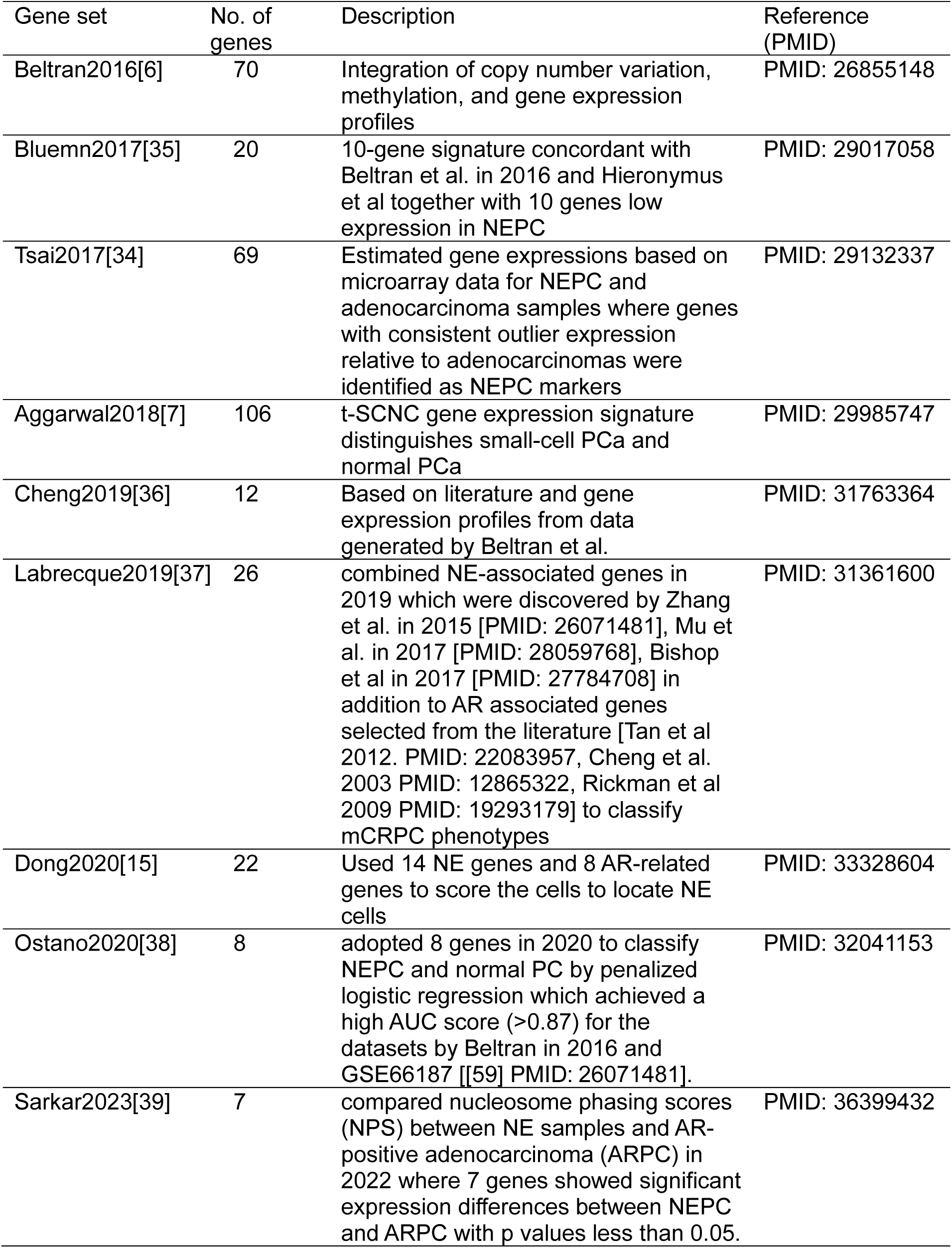
NEPC Marker gene sets in selected literatures.

**Supplemental Figure 1.**
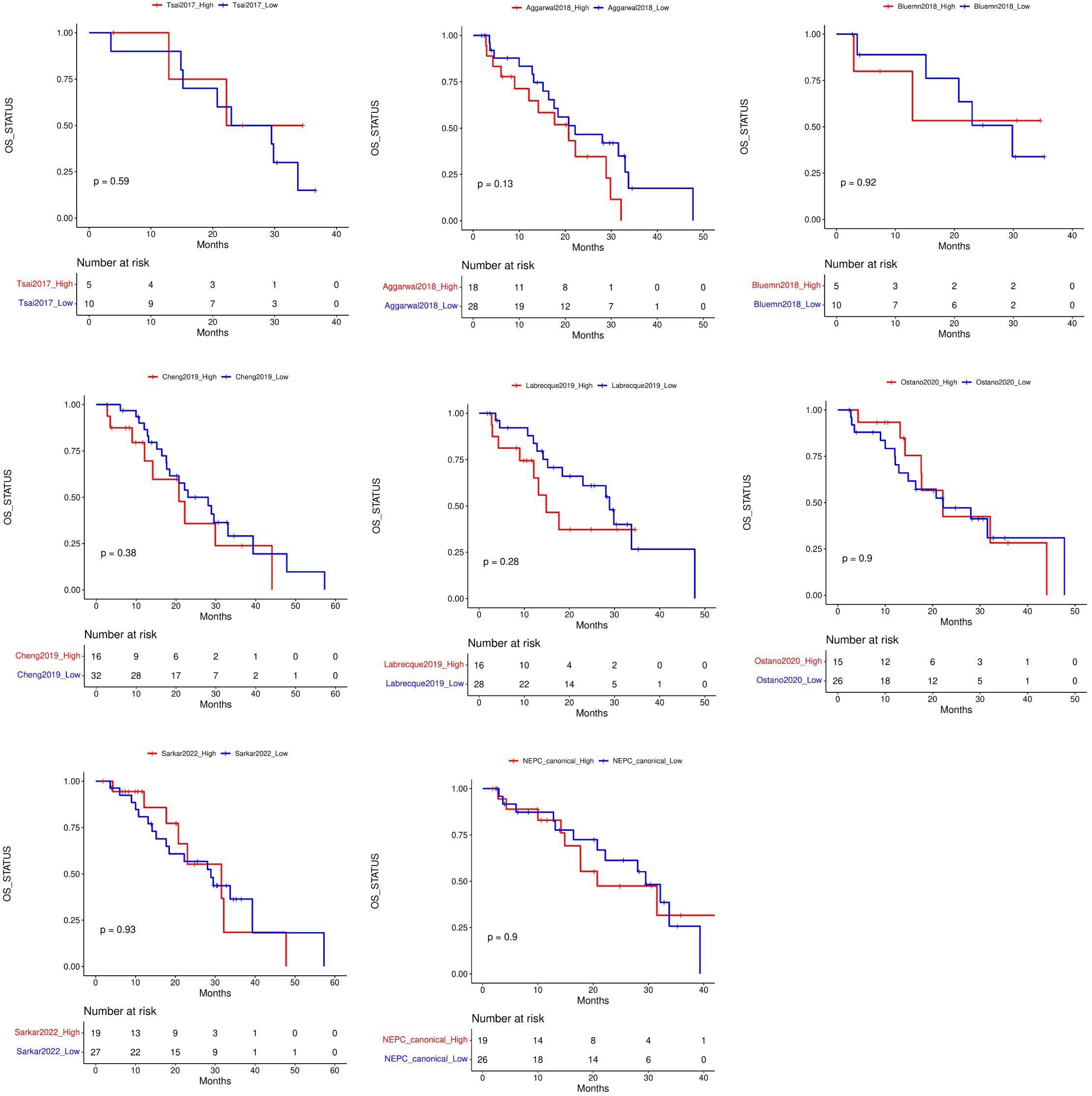
Survival plot of different gene sets on the dataset of PRAD_SU2C_2019.

**Supplemental Figure 2.**
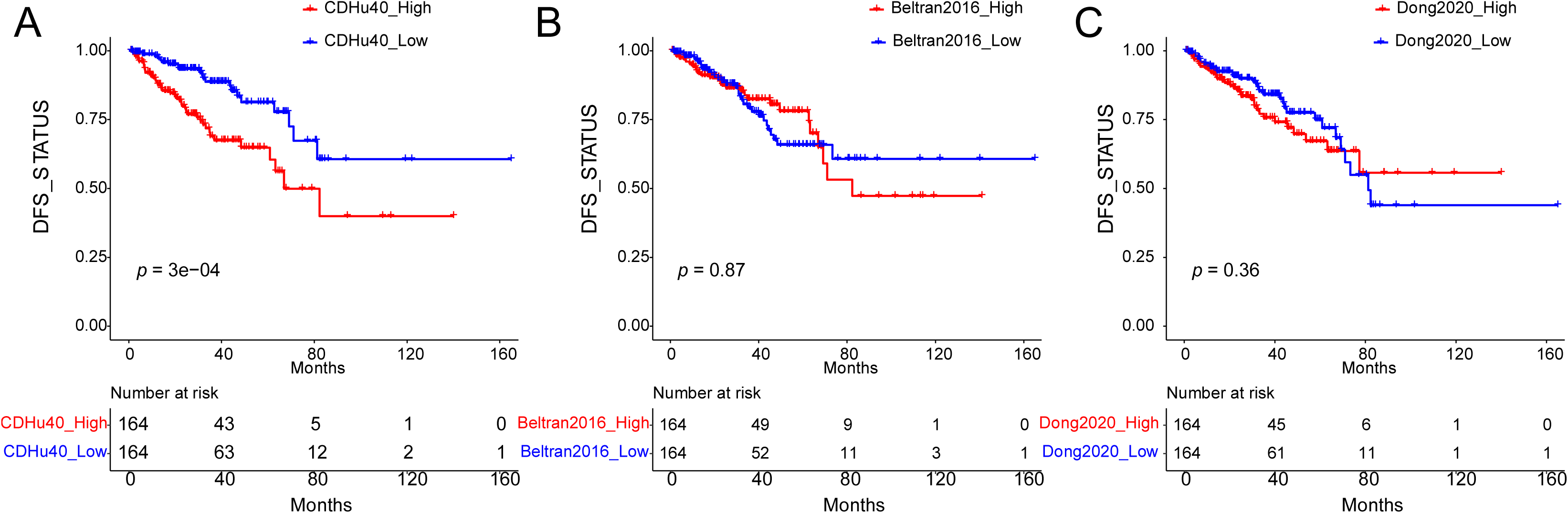
Survival difference between higher and lower scores on PRAD-TCGA samples identified by (A) CDHu40, (B) Beltran2016, and (C) Dong2020.

